# NEAR: Neural Embeddings for Amino acid Relationships

**DOI:** 10.1101/2024.01.25.577287

**Authors:** Daniel Olson, Thomas Colligan, Daphne Demekas, Jack W. Roddy, Ken Youens-Clark, Travis J. Wheeler

**Affiliations:** Department of Computer Science, University of Montana, Montana, USA; College of Pharmacy, University of Arizona, Arizona, USA

**Keywords:** Neural embedding, Protein sequence annotation, Contrastive Learning, Representation Learning

## Abstract

Protein language models (PLMs) have recently demonstrated potential to supplant classical protein database search methods based on sequence alignment, but are slower than common alignment-based tools and appear to be prone to a high rate of false labeling. Here, we present NEAR, a method based on neural representation learning that is designed to improve both speed and accuracy of search for likely homologs in a large protein sequence database. NEAR’s ResNet embedding model is trained using contrastive learning guided by trusted sequence alignments. It computes per-residue embeddings for target and query protein sequences, and identifies alignment candidates with a pipeline consisting of residue-level k-NN search and a simple neighbor aggregation scheme. Tests on a benchmark consisting of trusted remote homologs and randomly shuffled decoy sequences reveal that NEAR substantially improves accuracy relative to state-of-the-art PLMs, with lower memory requirements and faster embedding and search speed. While these results suggest that the NEAR model may be useful for standalone homology detection with increased sensitivity over standard alignment-based methods, in this manuscript we focus on a more straightforward analysis of the model’s value as a high-speed pre-filter for sensitive annotation. In that context, NEAR is at least 5x faster than the pre-filter currently used in the widely-used profile hidden Markov model (pHMM) search tool `HMMER3`, and also outperforms the pre-filter used in our fast pHMM tool, `nail`.

## Introduction

The ease and low cost of DNA sequencing creates unprecedented opportunity to catalog and understand the genomic diversity of life on Earth. One impact of the ongoing data deluge is that computational tools for sequence annotation should be designed with massive-scale search in mind [1, 2, 3]. Another is that new sequence data sets often include proteins that defy current annotation efforts – they are either entirely novel, or have diverged so far from their ancestral sequence that current methods are unable to detect their homology to already known sequence. The challenge is particularly great when annotating metagenomic datasets as they often reach terabases in size and large fractions (and in many cases, the majority) of encoded proteins may go unannotated [4, 5, 6] due to a combination of novelty, diversity, and sequencing error.

To meet these challenges, researchers continue to make advances to annotation methods along classical avenues of algorithm development and statistical model design [7, 8, 9, 10, 11, 12, 13]. Meanwhile, new strategies for annotation have gained traction, fueled by representation learning strategies [14] borrowed from the natural language processing (NLP) literature. NLP representation strategies have advanced rapidly from word2vec style [15] representation, to attention-based approaches [16] including BERT (Bidirectional Encoder Representations from Transformers [17]), T5 (Text-to-Text Transfer Transformer [18]), and GPT (Generative Pretrained Transformer [19]). Biosequence analogs to these methods [20, 21, 22, 23] learn representations of residues or sequences by employing standard unsupervised masked language model training. In the neural representation framework for proteins, a neural network computes a vector representation for a sequence, so that each sequence is embedded in a high dimensional representation space. A well-behaving protein language model (PLM) will embed a pair of sequences near each other in high-dimensional vector space if they share similar properties and will place dissimilar pairs far apart. Such models can be applied to sequence annotation with limited fine tuning, and have demonstrated tantalizing potential.

In a recent application of PLMs to detection of sequence relationships, Elnaggar et al. [22] showed that neural representation methods appear to identify some homologs of query proteins that are not found using sequence alignment methods. Importantly, the authors also showed that vector distances computed using their best-performing model struggled to distinguish decoys from true positives, so that the model was more useful when its top matches were re-scored by computing sequence alignments – in other words, it appears to be preferable to treat the model as a filter for more expensive sequence alignment (resulting in a loss in overall recall relative to `MMseqs2`).

In this work, we develop an alternate approach to learning and applying representations that enable fast and accurate recognition of relationships between proteins. We introduce a simple architecture and training method that computes a context-informed embedding of individual residues such that two residues will be close in embedding space if they are likely to be placed in the same column of a trusted pairwise sequence alignment. Our implementation (NEAR – Neural Embeddings for Amino acid Relationships) can be used to search a target data set T of proteins as follows: (1) for each sequence t of T, NEAR computes a high dimensional embedding vector for each residue in t and places that vector in a vector database index I; then (2) given a query sequence q, NEAR computes embeddings for each residue of q, searches I for near neighbors of each query residue, then identifies sequences from T that share many near neighbors with q.

For a new method to claim superior sensitivity over prior approaches (i.e. the ability to find previously-unrecognized relationships between proteins), it is important to first show that the new method at least matches the accuracy of prior methods (is able to recover known relationships without asserting verifiably false relationships). For this reason, we limit evaluation of NEAR and other tools to the recovery of trusted relationships, casting the problem as one of pre-filtering for alignment-based sequence homology tools. Future analyses will explore the more complex issue of annotation novelty. For a filter to be effective in a pipeline that seeks to find homologs of a query sequence q in a target set T or candidate proteins, it must identify a small subset of T that retains all known matches while removing nearly all decoy relationships, and it must do so quickly. We discuss these issues below.

### Search with high sensitivity and low decoy match rate

Because the most valuable filter is one that will enable maximally-sensitive downstream processing, we approach methods development and evaluation in the context of highly-sensitive annotation with profile hidden Markov models (pHMMs [24, 25, 12]), as employed in `HMMER` [26] and our `nail` [27]. Profile HMMs show greater sensitivity [28, 13, 7] than other homology search methods such as `BLAST` [29], LAST [30], and `MMseqs2` [7]. The sensitivity of pHMMs is due to a combination of (i) position specific scores [31] learned from sequence family members and (ii) implementation of the Forward algorithm [32, 24], which sums the probabilities of all possible alignments. The Forward algorithm is responsible for much of HMMER’s sensitivity gains, but is computationally expensive. `HMMER3` introduced a pipeline in which most candidates are never subjected to the most computationally expensive analysis, thanks in large part to a stage called MSV that performs highly-optimized ungapped sequence alignment to identify promising seeds. By default, `HMMER3` ’s MSV stage filters away all but *∼* 2% of decoy (non-homologous) sequences, while retaining essentially all pairs that would pass a full Forward analysis. In common usage, the MSV filter accounts for *∼*70% of `HMMER3` ’s run time [33]. Meanwhile, our high-speed pHMM tool, `nail`, employs a reparameterized `MMseqs2` as pre-filter; this approach is fast but lossy, and the primary cause of the sensitivity gap between `nail` and HMMER3 [27]. NEAR is designed to fulfill the role played by MSV in the `HMMER3` pipeline and `MMseqs2` in the `nail` pipeline: given a large set T of target protein sequences and a query protein sequence, rapidly identify a small subset of target sequences that are expected to produce high Forward scores, so that a large majority of the unrelated sequences in T can be ignored (filtered) without performing expensive sequence alignment. Ideally, a neural filter such as NEAR will filter as effectively as `HMMER3`’s MSV filter, ranking all true matches with score greater than the vast majority of matches to decoys.

### Search speed

In addition to at least matching the filtering efficacy of `HMMER3`’s MSV filter, an ideal filter will improve on the filtration speed. To achieve acceptable speed, a neural filter must first compute embeddings quickly; NEAR’s model is smaller than state-of-art PLMs (at least 60-fold fewer parameters; see “Model Architecture”), and thus computes residue-level embeddings much more quickly. A neural filter must also store the set of target embeddings in an efficient data structure that supports rapid identification of k nearest neighbors without computing distances for all neighbors. Fast approximate near neighbor search in high volume and high dimensional data is a highly researched area [34, 35, 36, 37, 38]. NEAR uses the Facebook AI Similarity Search (FAISS) library [39].

### Manuscript focus and software availability

In the sections below, we provide a description of methods for model design, training, and searching. We then demonstrate that NEAR’s similarity calculations enable recovery of true matches with greater filtering efficacy than other neural embedding strategies, and is faster than both PLM-based filters and HMMER’s MSV filter. NEAR is available under an open license at https://github.com/TravisWheelerLab/NEAR.

## Methods

### Model architecture

NEAR’s embedding model is implemented as a 1D Residual Convolutional Neural Network (ResNet) [40]. Each of NEAR’s residual blocks performs the following computation:

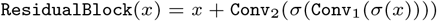

`Conv`_1_ and `Conv`_2_ are learned 1D-convolutions with the same hyper-parameters, and σ is an activation function. There is no weight-sharing between convolutional layers or residual blocks. NEAR’s default model hyper-parameters can be seen in Table

1. The total number of parameters for the NEAR ResNet is *∼*7M parameters, in contrast to *∼*3B for the tested variant of ProtTransT5, *∼*600M for ESM C’s largest open model, and *∼*420M for ProtBERT.

### Model Training

NEAR’s neural network transforms a sequence of amino acids, S = {S_1_, S_2_, …, S_n_} into a sequence of high dimensional vectors,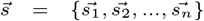. Each vector 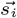 represents the corresponding residue, S_i_, in a manner that reflects the residue’s surrounding context. NEAR learns to embed sequences so that residues that align to one another within trusted alignments will be embedded as vectors that have a large dot product, while unaligned residues will typically have small dot products.

NEAR’s training procedure (see Figure 1), takes a pair of homologous sequences, A and B, and generates a target matrix, T, based on the `phmmer`-alignment of A and B such that T_ij_ = 1 when A_i_ aligns with B_j_, and is 0 everywhere else (Figure 1A right). Separately, NEAR’s neural network is used to embed the residues of A and B into high dimensional vectors,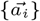 and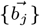, for all 0 < i *≤* |A| and 0 < j *≤* |B|. Importantly, A and B are embedded independently of one another; NEAR’s neural network does not use the embedding of one sequence to inform its embedding of the other sequence. The embeddings are used to create a matrix, D, where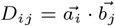 (Figure 1A right).

**Fig. 1.**
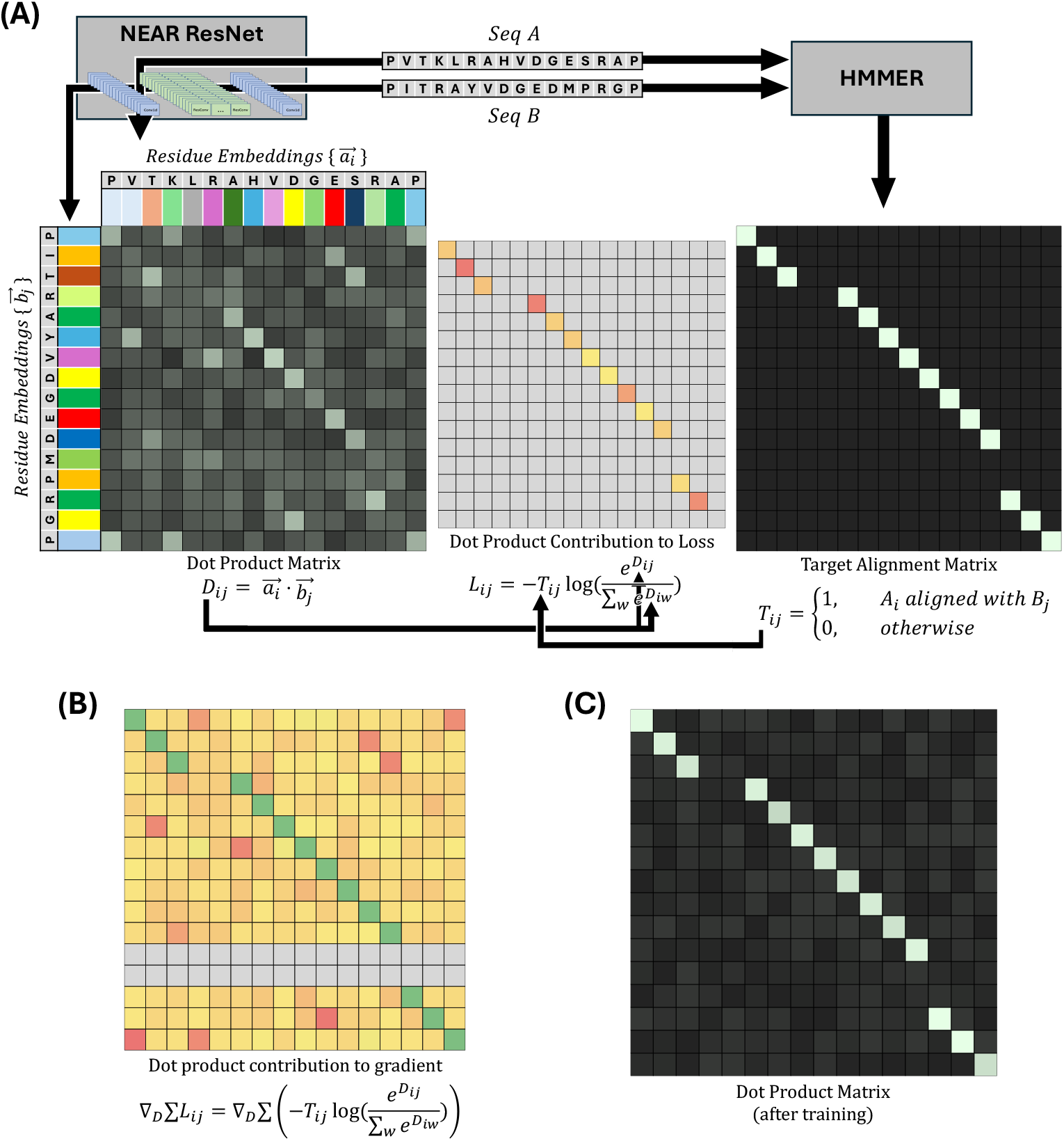
**(A)** shows a cartoon description of NEAR’s training procedure. Pairs of homologous sequences (A and B) are embedded with NEAR to produce residue-embeddings,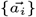 and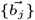, represented as colored rectangles. The dot products of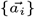 and 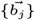form a matrix, D. An untrained neural network will produce residue-embeddings that poorly represent the alignment expectation of those residues, resulting in noisy and uninformative dot products. NEAR’s neural network is trained by applying an N-pair loss to D using a target matrix, T, generated from the `phmmer` alignment of A and B. **(B)** shows the contribution of individual dot products on the loss gradient. Although N-Pair loss produces a sparse loss signal (shown in loss matrix, L), the gradient for N-pair loss is dense, i.e the majority of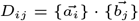dot products have some effect on the gradient of NEAR’s model parameters. To minimize the N-pair loss, L, the softmax,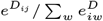, must be maximized wherever T_ij_ = 1, resulting in gradients that encourage D_ij_ to be large wherever T_ij_ = 1 (maximizing the softmax numerator) and discourage D_ij_ from being large wherever T_ij_ = 0 (minimizing the softmax denominator). **(C)** shows a cartoon dot product matrix of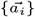and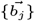 when embedding A and B with a fully trained ResNet.

NEAR uses a loss function composed of three elements: (1) An N-Pair loss [42] that drives vector pairs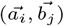 in similar directions when residues (A_i_, B_j_) share an alignment column. (2) A masking factor, R, that reduces problems caused by repetitive and low-complexity sequence regions. (3) An L_2_ regularization term (controlled by a scaling factor, γ) that prevents embeddings from becoming unnecessarily large.

N-Pair loss is calculated by performing a summation of negative log softmax values for all aligned (i, j) residue pairs. Although the loss is only explicitly applied to aligned vector pairs (i.e (i, j) pairs where T_ij_ = 1), unaligned residues still propagate meaningful gradient (Figure 1B): when T_ij_ = 1, the loss is improved not only by increasing the softmax numerator,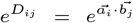, but also by decreasing the softmax denominator,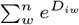. In this formulation, high similarity between unaligned vectors 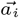 and 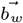 will increase the denominator, causing a reduction in the softmax value of the aligned vectors, 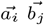. As a result, minimization of N-Pair loss not only drives aligned embeddings in similar directions, but also pushes unaligned embeddings in orthogonal directions.

Repetitive and low complexity sequence regions pose a challenge to similarity-based homology search methods. Typically, two protein sequences that have high sequence-similarity to one another can be confidentially annotated as homologous. However, sequences that share similar composition bias/repetitive content may have high similarity scores to one another regardless of whether or not the sequences are evolutionarily related to one another. NEAR uses a repeat masking strategy similar to those used in classical sequence alignment [43, 44] to account for problems caused by repetitive sequence both during search and also during training: when learning to embed a sequence S during training, NEAR uses the software `tantan` [43] to create a mask *R*^*s*^, assigning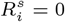when S_i_ is labeled as repetitive by `tantan`, and 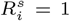 everywhere else. This mask is used to silence loss signals produced in repetitive/baised regions on training.

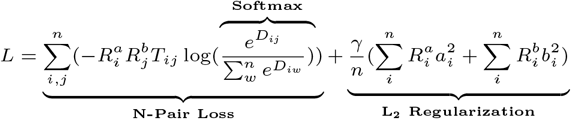

#### Training hyper-parameters

Hyper-parameters used to train NEAR can be seen in Table 2. During training, NEAR utilizes the AdamW optimizer [45].

**Table 1.** Model hyper parameters.

| Hyper-parameter | Value |
| --- | --- |
| Dimensionality | 256 |
| Residual blocks | 8 |
| Kernel size | 7 |
| Activation function, $\sigma$ | ELU[41] |

**Table 2.**
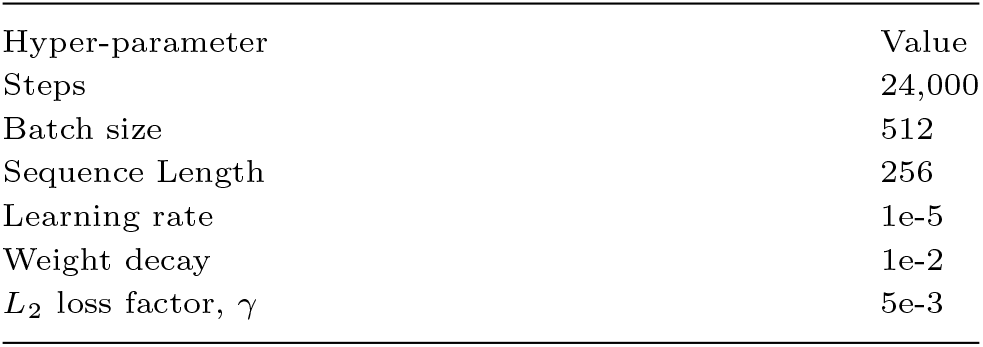
Training hyper parameters.

### Sequence search

#### Vectors for finding similar residues

Search against a set of target sequences, T, begins with creation of a search index over those sequences. NEAR computes residue-embeddings for the sequences in T, and places the resulting vectors in a search index I built using the FAISS library [39] (using the `IVF5000,PQ32` search index).

With this search index I in hand, NEAR computes residue vectors for the query protein; then for each residue vector, it employs the FAISS library to identify similar vectors in I. This nearest neighbor search results in a list of (default) 150 residue *matches* for each query residue embedding. Two d-dimensional random vectors will occasionally produce moderately-sized cosine similarities by chance. We reduce the effect of cosine similarity noise by subtracting a noise gate term,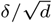.

The standard deviation of cosine similarities for d-dimensional vectors drawn from 𝒩(0, *I*) grows inversely proportional to 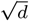, and so scaling the noise gate parameter, *δ*, by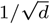 allows *δ* to have consistent effect across different embedding dimensionalities. By default NEAR uses *δ* = 3, filtering out an expected 99.9% of cosine similarity noise. With this in mind, the strength of a match is measured as the cosine similarity between the query and target embeddings, adjusted by the noise gate. The score of residue pair (A_i_, B_j_) is:

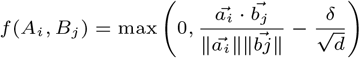

NEAR avoids false-positive search results caused by compositional bias and tandem repeats using `tantan` [43] to identify residues in repetitive/biased regions and then excluding the associated residue embeddings from the search pipeline (this is similar to the masking strategy used in sequence alignment methods [46]).

#### Score accumulation

After using FAISS to find similar query and target *residue* embeddings, NEAR uses a simple mechanism to accumulate the support for sequence-level similarity. Let M_A,B_ be the set of matching residue pairs found during FAISS search for some query sequence, A, and some target sequence, B. To estimate the overall similarity between A and B, NEAR combines all matching residue pairs like so:

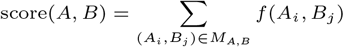

#### Datasets

We used Uniclust30 [47] to gather clusters of UniRef [48] sequences that were between 256 and 1024 residues in length. In this dataset, sequences in one cluster are <30% identical to sequences in any other cluster. Clusters with only one member were discarded, and 80% of the remaining clusters were used to construct a training set, while the remaining 20% were used to construct a sequence search benchmark to be used during evaluation. We will refer to the training clusters as **C**_train_ and the clusters used for benchmarking as **C**_eval_.

#### Training data

Sequence pairs and `HMMER3`-generated alignments are used during training to guide NEAR’s contrastive learning objective (see Figure 1 and related text). We created a collection of sequence pairs/alignments by randomly sampling sequence pairs from each cluster in **C**_train_ and then aligning those sequences pairs to one another using the HMMER command, `phmmer --max`, which performs sequence-to-sequence alignment using HMMER’s Forward algorithm. To ensure a broad training distribution, no cluster was allowed to contribute more than 25 sequence pairs. This process resulted in 6,190,084 sequence pairs and alignments that were used during training.

#### Evaluation data

The evaluation of NEAR (see Results) utilized three sets of protein sequences: **Q, T**^+^, and **T**^*−*^. **Q** and **T**^+^ were produced by randomly selecting 10,000 clusters from **C**_eval_, each cluster containing between 10 and 50 sequences, and then randomly sampling (without replacement) sequences from those clusters. From this procedure we curated a collection of 10,000 protein sequences, **Q**, to be used as homology search queries, and a set of 20,000 protein sequences, **T**^+^, to be used as targets. We also constructed a set of decoy sequences, **T**^*−*^, that have no meaningful biological relationship to sequences in **Q** or **T**^+^. **T**^*−*^ was generated by shuffling the contents of each sequence in **T**^+^.

Our evaluation also relies upon trusted **Q** to **T**^+^ sequence similarities. These similarities were generated by first repeat-masking sequences in **Q** and **T**^+^ using `tantan` -w 16 -p [43], and then aligning repeat-masked **Q** and **T**^+^ sequences using `phmmer --max`.

### Evaluation against other search tools

In our evaluation of NEAR (see 3) we compare NEAR against classical homology search pre-filter tools and also other deep-learning based approaches. More specifically, we compare NEAR against `MMSeqs2`’s pre-filter [7], `HMMER3`’s pre-filter (MSV) [26], ProtTransT5 [22] (`prot t5 xl half uniref50-enc)`, ESM C [**?**] (`ESM C 600M`), ProtBert [21], and TM-Vec [49].

We compare the filtering efficacy of NEAR to that of classical filtering algorithms in `MMSeqs2` and `HMMER3`. `MMSeqs2` uses an extremely fast (but less sensitive) kmer-hashing approach to find potential sequence-alignment candidates. `HMMER3` uses a sensitive (but slower) gapless alignment algorithm called MSV (Multi-Segment ungapped Viterbi) [26].

ProtTransT5, ESM, and ProtBert are *protein language models* (aka PLMs) that are used to create embeddings for protein sequences and residues. The embeddings produced by these PLM’s are more general-purpose than the embeddings produced by NEAR and have been used to infer protein homology [50], structure [51], and function [52]. To use PLMs for homology search, we generated embeddings of **Q** and **T** using the PLMs, and then used those embeddings in the FAISS sequence similarity pipeline search akin to the usage for NEAR. We measure PLM performance when using sequence-level (residue-averaged) embeddings (which is common practice), and also when using PLM residue embeddings with NEAR’s sequence search pipeline.

TM-Vec is one of several methods (see [53, 50, 54]) that use PLMs to create protein sequence embeddings that are then fed into a small neural network to produce embeddings specifically for protein similarity search; TM-Vec’s neural network was trained to predict TM-score [55]. All TM-Vec results were gathered using TM-Vec’s built in vector search pipeline.

While NEAR (by default) produces 256 dimensional embeddings, ProtTransT5 and ProtBert both produce 1024 dimensional embeddings, and ESM produces 1152 dimensional embeddings. PLM residue embeddings of **T**’s 40,000 sequences were too large to fit into GPU memory for FAISS search. For this reason, we used a reverse search scheme when performing FAISS search with PLM embeddings, wherein we built a FAISS search index using the 10,000 query sequences, **Q**, and searched **T** against that index. This reversed-search approach is expected to slightly improve maximum recall relative to a standard query-to-target search, and this is observed when a similar search orientation was performed with NEAR. For all neural embedding tests performed with this inverted search orientation, the name of the tool is supplemented with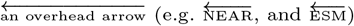.

## Results

Sequence search pre-filters aim to rapidly reduce the set of all query-target pairs down to only those that are likely to score well in downstream analysis (sequence alignment). We evaluated the competing tools (`MMSeqs2`, `HMMER3`’s MSV, ProtTransT5, ESM, ProtBert, and TM-Vec) by using each tool to perform sequence search between **Q** and **T**, measuring performance as the percentage of homologous **Q**-**T**^+^ sequence pairs recovered at varying levels of decoy filtration.

One challenge in fair evaluation of homology search tools (and homology search pre-filters) is that it is difficult or even impossible to know with certainty all homology relationships within a diverse collection of protein sequences. Although we can infer with high confidence that highly similar sequences are homologous, we cannot rule out the possibility that two less-similar sequences may be still evolutionarily related. We try to address this challenge by examining results using two different filtration measurements: Filtration of nonhomologous **Q**-**T**^*−*^ sequence pairs and filtration of low-similarity **Q**-**T**^+^ pairs.

### Filtration of nonhomologous decoy sequence pairs

Figure 2 compares pre-filter recall of high similarity sequence pairs against filtration of *nonhomologous sequence pairs*, i.e sequence pairs containing one true protein sequence and one decoy (shuffled) sequence, **Q**-**T**^*−*^. An advantage of only measuring filtration of **Q**-**T**^*−*^ pairs is that tools will not be penalized for finding biologically meaningful relationships between protein sequences that are missed by traditional alignment-based homology search approaches. The set of shuffled sequences, **T**^*−*^, are expected to have no meaningful structure, function, or similarity to sequences in **Q**, and so pre-filter tools should filter out all **Q**-**T**^*−*^ sequence pairs. One disadvantage to this approach is that it may overestimate pre-filter performance; it is generally easier to differentiate between biological protein sequences and random protein sequences than it is to differentiate between biological protein sequences and unrelated biological protein sequences.

**Fig. 2.**
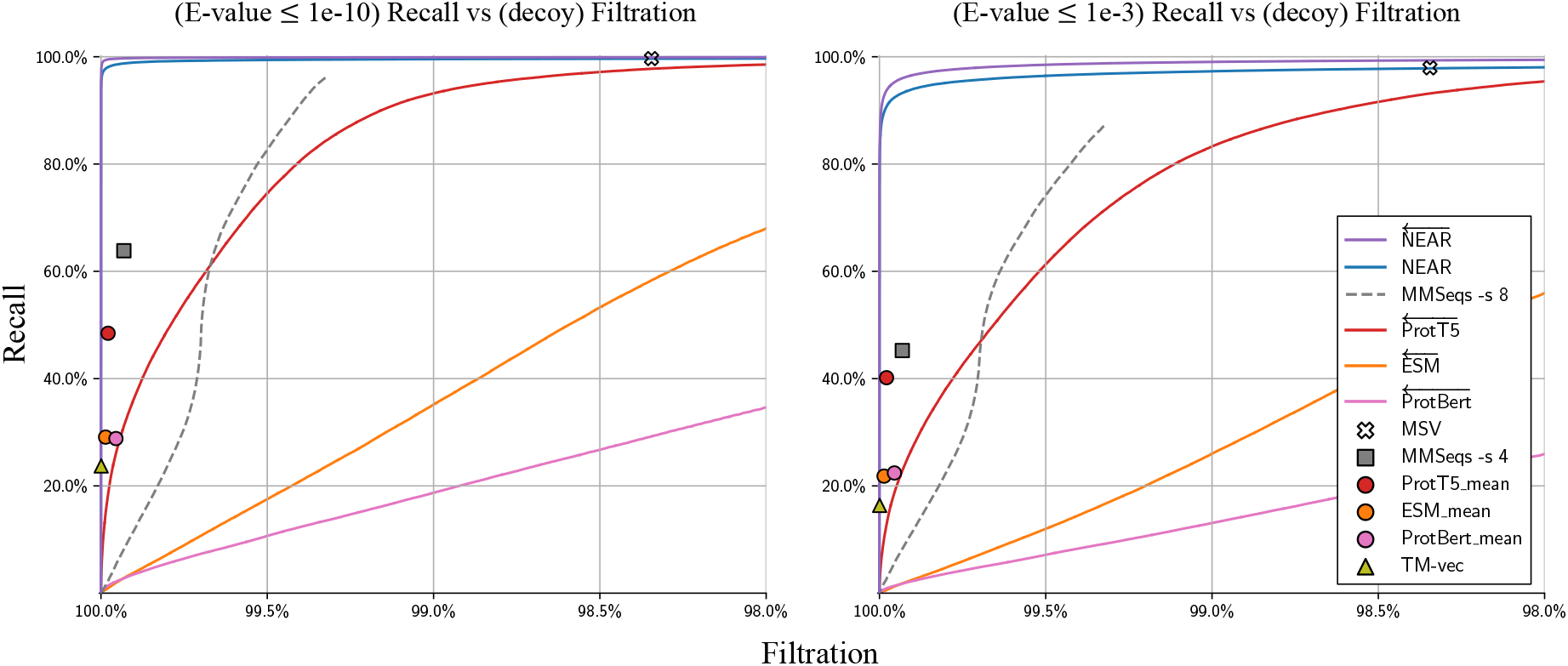
Recall of high-similarity sequence pairs measured against filtration of sequence pairs containing shuffled decoy sequences (i.e **Q-T**™ sequence pairs). Curves were produced by sorting **Q**-**T** hits according to pre-filter score; each point on a curve can be interpreted as the recall/filtration when using a particular pre-filter score threshold. Methods that produced high filtration rates regardless of score-threshold are represented as points instead of curves. For all neural embedding methods, the curves present recall/filtration results for the approach of using the model to produce per-residue embeddings, then accumulating scores as in NEAR. For each PLM tool, an additional result is plotted as a colored shape; this shows recall when embedding only a single vector per sequence (computed as the mean of the individual residue vectors). Our vector search for single sequence embeddings can capture at most 150 nearest neighbor target sequence embeddings for each query sequence embedding. This is in line with prior applications and sufficient to achieve 100% recall, but results in a high base filtration rate; as a result, embedding search methods that use a single sequence-embedding are presented as points and not curves. The left plot measures recall of **Q**-**T**^+^ sequence pairs that have `phmmer` E-values ≤ 1e-10, and the right plot measures recall of **Q**-**T**^+^ sequence pairs with `phmmer` E-values ≤ 1e-3.

Figure 2 shows recall (y-axis) as a function of filtration for all tools on two datasets. On the left, the set of true matches (which are used to measure recall) is defined as the collection of pairs from **Q**-**T**^+^ in which `phmmer --max` produces an E-value *≤* 10^*−*10^ – these are strong-scoring `phmmer` matches that should not be missed by any homology detection tool.

On the right, the stringency for *true* matches is reduced to E-value *≤* 10^*−*3^; this set includes some lower confidence (or possibly spurious) matches, and it is not surprising for pre-filters to struggle to assign high scores to all pairs. Under both similarity thresholds,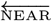 shows the greatest accuracy followed shortly by `HMMER3`’s MSV. Although PLM’s are often thought to encode a rich understanding of protein sequences, they provided surprisingly lackluster performance in differentiating high-similarity sequence pairs from the decoy **Q**-**T**^*−*^ sequence pairs.

#### Filtration of low similarity sequence pairs

In the previous test, decoys were defined by providing sequence (**T**^*−*^) that are shuffled, and therefore expected to have no retained protein structure. Figure 3 assesses filtration using an alternate set of decoys based on the real (un-shuffled) proteins in **T**^+^: `phmmer` --max was used to force an alignment between all pairs in **Q**-**T**^+^; pairs with E-value *≥* 10 were identified as having no alignment-supported relationship, and established as *decoys*. By only measuring recall and filtration of **Q**-**T**^+^ sequence pairs (i.e purely biological protein sequence pairs) these results may serve as a more realistic indicator of pre-filter performance. Although low-similarity sequence pairs are not guaranteed to be unrelated to one another, (a)it is safe to filter the pairs, since these matches would not be reported under typical annotation parameters, and (b)it is reasonable to expect that the majority of sequence pairs with E-value *≥* 10 are nonhomologous. It is useful for a pre-filter to remove most of these candidates while retaining recall on low E-value pairs. For example, at a score threshold that filters 80% of very-low-scoring matches to real protein sequences (**Q**-**T**^+^ with E-value >10), NEAR yielded the greatest recall, demonstrating NEAR’s utility in quickly removing most unnecessary comparisons before downstream analysis.

**Fig. 3.**
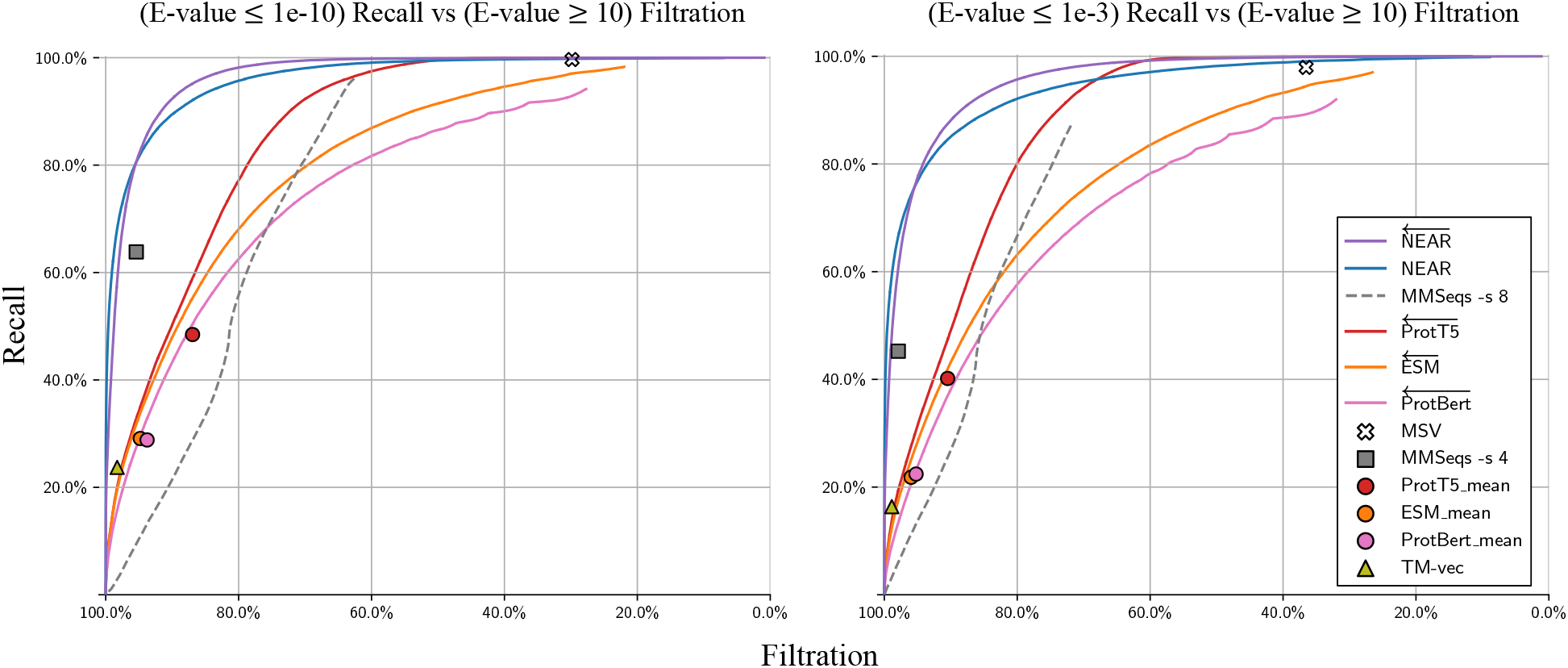
Recall of high-similarity sequence pairs measured against filtration of low-similarity **Q**-**T**^+^ sequence pairs. We define low-similarity sequence pairs as those with E-values ≥ 10. The left plot measures recall of **Q**-**T**^+^ sequence pairs with `phmmer` E-values ≤ 1e-10, and the right plot measures recall of **Q**-**T**^+^ sequence pairs with `phmmer` E-values ≤ 1e-3.

### Compute time

Table 3 reports the time required to embed sequences, build search indices, and perform a **Q**-**T** search. All computations were performed on a g3xl Jetstream2 [56, 57] instance using an AMD EPYC-Milan Processor and A100-SXM4-40GB GPU. Timings were collected using a single GPU (for embedding-search methods) and using a single CPU (for `MMSeqs2` and MSV).

**Table 3.**
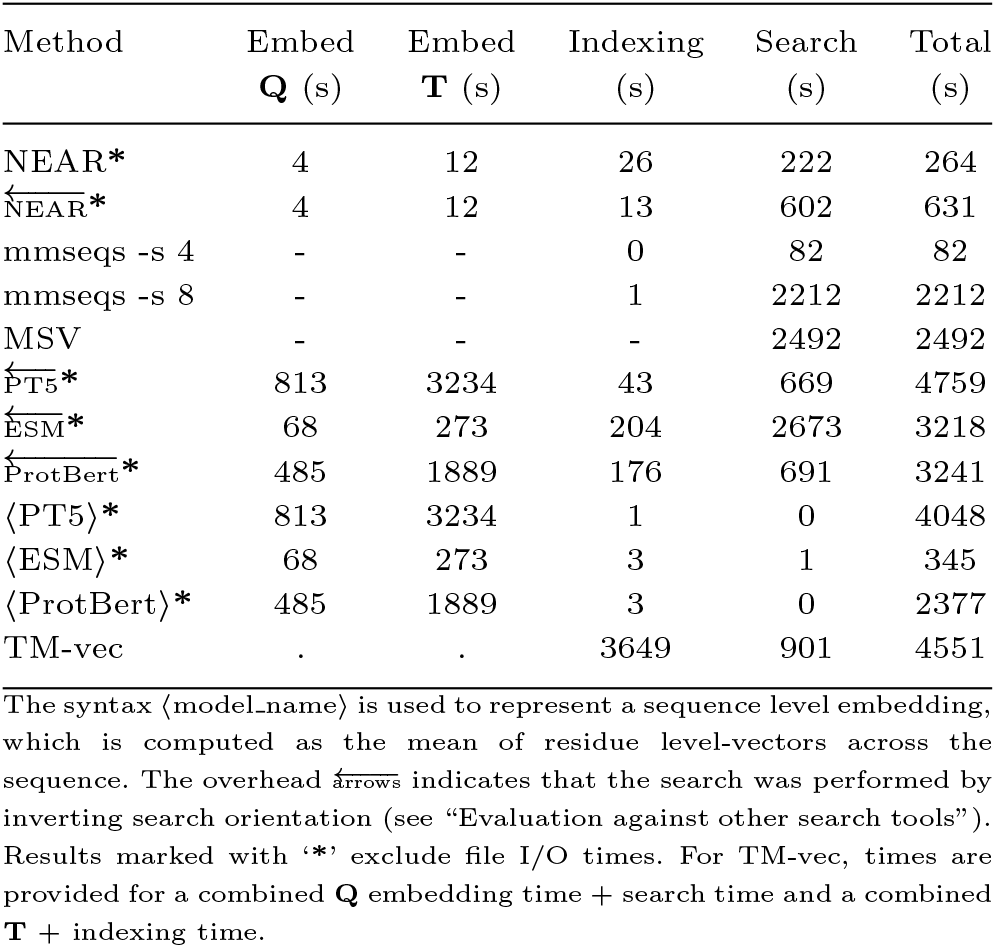
Compute times for embedding, indexing, and searching 10,000 query sequences, **Q**, against 40,000 target sequences, **T**.

While the Jetstream2 system is an excellent resource for model training, the instances are virtualized and file access across a distributed file system leads to an expensive file I/O overhead. When evaluating neural embedding methods with our FAISS search pipeline we evaluated embedding and search separately, resulting in an inflated runtime due to the high file I/O cost. To better estimate the end-to-end runtime of NEAR, we have excluded file I/O time from methods that use our FAISS search pipeline (NEAR, PT5, ESM, and ProtBert).

## Discussion

Here, we have described NEAR, a neural network and search pipeline designed to act as a fast and sensitive homology search pre-filter. NEAR builds on existing concepts in Representation learning by implementing a simple ResNet architecture and a novel training regimen that enables faster and more accurate search than state of art transformer-based protein language models (such as ProtTransT5, ESM, ProtBERT). NEAR’s model requires < 2% of the parameters of those PLMs, and utilizes a lower dimensionality representation space; combined, these features are responsible for NEAR’s speed and memory improvements. NEAR is also competitive with algorithmic approaches to alignment filtering, achieving greater accuracy and greater speed than HMMER’s pre-filter, while improving on `MMseqs2`’s (high-sensitivity) filtering accuracy with comparable speed. These results collectively demonstrate the efficacy of NEAR in accurately ranking hits.

### Interpretation of embedding vectors

General purpose embeddings produced by PLMs encode an impressive breadth of complex information. PLM embeddings can be used to predict protein structure and protein function. Additionally, they can be searched or clustered to reveal evolutionary and functional relationships missed by other tools. A downside to the generality provided by PLM embeddings it is difficult to understand the totality of information encoded within a PLM embedding; as a result, it is difficult to interpret the meaning and significance of two PLM embeddings being similar to one another.

The representational power of NEAR’s neural network is extremely limited relative to the PLMs tested in this paper.

The inductive biases of NEAR’s ResNet architecture restrain what NEAR’s model *can* learn; NEAR can only learn to create residue-embeddings based on local sequence context. The specificity of NEAR’s training task further restrict what the model *does* learn; NEAR learns to embed residues so that the similarity of two residue-embeddings corresponds to how well the corresponding residues are expected to align with one another (which itself likely corresponds to how similar the sequence regions are to each other). High similarity between only two residue embeddings does not indicate that the corresponding sequences will align well to one another; instead, it indicates that there is at least one strong *alignment seed* where the sequences are likely to locally align to one another. NEAR calculates a similarity score between two sequences by summing (noise gated) cosine-similarities for all similar embeddings between the two sequences; this sum effectively represents the total number of good alignment seeds between the two sequences.

### Role of representations in homology detection

Though NEAR shows promise as a fast and accurate filter for pHMM search, we have not created an integrated tool by connecting NEAR with pHMM software; in the future, we intend to explore the use of NEAR with our pHMM tool, `nail` [27]. Here, we have focused on designing an effective model architecture and training strategy, and also performing experiments to explore NEAR’s viability as a pre-filter. Counter-intuitively, NEAR works well as a pre-filter not by compressing data, but by expanding data. NEAR transforms simple sequence strings into large and complex sequences of high dimensional vectors, enabling us to harness the wealth of research and software available in the field of vector search. An important concern regarding NEAR’s utility as an alignment filter is that by representing residues as high dimensional vectors, NEAR has limited utility on large datasets; precisely where fast filters are most important. Using a quantized search index allows for a modest reduction in memory and compute costs, but a much greater reduction is necessary for NEAR to be a viable pre-filter method for large sequence databases (note: *all* residue-level PLM embedding approaches suffer the same challenges). Advances to methods for sketching or alternative sparse vector sequence representation, perhaps based on prediction of which residues are important for search, will be vital for improved scalability. Finally: we anticipate that embedding similarity values may be useful as a source of position-specific scores for direct sequence alignment [58, 50].

## Acknowledgments

We thank Tim Anderson and Genevieve Krause for helpful discussions during development of software and benchmarks. We also gratefully acknowledge the computational resources and expert administration provided by the University of Montana’s Griz Shared Computing Cluster (GSCC), CyVerse’s External Collaborative Partnership program (with special thanks to Tyson Swetnam), and the high performance computing (HPC) resources supported by the University of Arizona TRIF, UITS, and Research, Innovation, and Impact (RII) and maintained by the UArizona Research Technologies department. Compute was also supported through Jetstream2 [56] g3xl compute nodes provided by allocation CIS240916 from the Advanced Cyberinfrastructure Coordination Ecosystem: Services & Support (ACCESS) program[57], which is supported by National Science Foundation grants #2138259, #2138286, #2138307, #2137603, and #2138296. This work was supported by NIH NIGMS R01GM132600, by DOE BER DE-SC0021216.

## References

1. C Titus Brown and Luiz Irber. sourmash: a library for MinHash sketching of DNA. Journal of open source software, 1(5):27, 2016.

2. Sewon Lee, Gyuri Kim, Eli Levy Karin, Milot Mirdita, Sukhwan Park, Rayan Chikhi, Artem Babaian, Andriy Kryshtafovych, and Martin Steinegger. Petascale homology search for structure prediction. bioRxiv, 2023.

3. Robert C Edgar, Brie Taylor, Victor Lin, Tomer Altman, Pierre Barbera, Dmitry Meleshko, Dan Lohr, Gherman Novakovsky, Benjamin Buchfink, Basem Al-Shayeb, et al. Petabase-scale sequence alignment catalyses viral discovery. Nature, 602(7895):142–147, 2022.

4. Quentin Carradec, Eric Pelletier, Corinne Da Silva, Adriana Alberti, Yoann Seeleuthner, Romain Blanc-Mathieu, Gipsi Lima-Mendez, Fabio Rocha, Leila Tirichine, Karine Labadie, et al. A global ocean atlas of eukaryotic genes. Nature communications, 9(1):373, 2018.

5. Eli Levy Karin, Milot Mirdita, and Johannes Söding. MetaEuk—sensitive, high-throughput gene discovery, and annotation for large-scale eukaryotic metagenomics. Microbiome, 8:1–15, 2020.

6. Sejal Modha, David L Robertson, Joseph Hughes, and Richard J Orton. Quantifying and cataloguing unknown sequences within human microbiomes. Msystems, 7(2):e01468–21, 2022.

7. Martin Steinegger and Johannes Söding. MMseqs2 enables sensitive protein sequence searching for the analysis of massive data sets. Nature biotechnology, 35(11):1026–1028, 2017.

8. Benjamin Buchfink, Klaus Reuter, and Hajk-Georg Drost. Sensitive protein alignments at tree-of-life scale using DIAMOND. Nature methods, 18(4):366–368, 2021.

9. Kristoffer Sahlin. Effective sequence similarity detection with strobemers. Genome research, 31(11):2080–2094, 2021.

10. Martin C Frith. A simple method for finding related sequences by adding probabilities of alternative alignments. Genome Research, 34(8):1165–1173, 2024.

11. Martin C Frith. A new repeat-masking method enables specific detection of homologous sequences. Nucleic acids research, 39(4):e23–e23, 2011.

12. Sean R. Eddy. Profile hidden Markov models. Bioinformatics, 14(9):755–763, 1998.

13. Genevieve R. Krause, Walt Shands, and Travis J. Wheeler. Sensitive and error-tolerant annotation of protein-coding DNA with BATH. bioRxiv, 2024.

14. Yoshua Bengio, Aaron Courville, and Pascal Vincent. Representation learning: A review and new perspectives. IEEE transactions on pattern analysis and machine intelligence, 35(8):1798–1828, 2013.

15. Tomas Mikolov, Kai Chen, Greg Corrado, and Jeffrey Dean. Efficient estimation of word representations in vector space. arXiv preprint, 2013.

16. Ashish Vaswani, Noam Shazeer, Niki Parmar, Jakob Uszkoreit, Llion Jones, Aidan N Gomez, Lukasz Kaiser, and Illia Polosukhin. Attention is all you need. Advances in neural information processing systems, 30, 2017.

17. Jacob Devlin, Ming-Wei Chang, Kenton Lee, and Kristina Toutanova. Bert: Pre-training of deep bidirectional transformers for language understanding. In Proceedings of the 2019 Conference of the North American Chapter of the Association for Computational Linguistics: Human Language Technologies, pages 4171–4186. Association for Computational Linguistics, 2019.

18. Colin Raffel, Noam Shazeer, Adam Roberts, Katherine Lee, Sharan Narang, Michael Matena, Yanqi Zhou, Wei Li, and Peter J Liu. Exploring the limits of transfer learning with a unified text-to-text transformer. The Journal of Machine Learning Research, 21(1):5485–5551, 2020.

19. Alec Radford, Karthik Narasimhan, Tim Salimans, Ilya Sutskever, et al. Improving language understanding by generative pre-training. 2018.

20. Dhananjay Kimothi, Akshay Soni, Pravesh Biyani, and James M Hogan. Distributed representations for biological sequence analysis. arXiv preprint, 2016.

21. Nadav Brandes, Dan Ofer, Yam Peleg, Nadav Rappoport, and Michal Linial. ProteinBERT: a universal deep-learning model of protein sequence and function. Bioinformatics, 38(8):2102–2110, 2022.

22. Ahmed Elnaggar, Michael Heinzinger, Christian Dallago, Ghalia Rihawi, Yu Wang, Llion Jones, Tom Gibbs, Tamas Feher, Christoph Angerer, Martin Steinegger, et al. ProtTrans: Toward understanding the language of life through self-supervised learning. IEEE transactions on pattern analysis and machine intelligence, 44(10):7112– 7127, 2021.

23. Alexander Rives, Joshua Meier, Tom Sercu, Siddharth Goyal, Zeming Lin, Jason Liu, Demi Guo, Myle Ott, C Lawrence Zitnick, Jerry Ma, et al. Biological structure and function emerge from scaling unsupervised learning to 250 million protein sequences. Proceedings of the National Academy of Sciences, 118(15):e2016239118, 2021.

24. Anders Krogh, Michael Brown, I Saira Mian, Kimmen Sjölander, and David Haussler. Hidden Markov models in computational biology: Applications to protein modeling. Journal of molecular biology, 235(5):1501–1531, 1994.

25. Richard Durbin, Sean R Eddy, Anders Krogh, and Graeme Mitchison. Biological sequence analysis: Probabilistic models of proteins and nucleic acids. Cambridge university press, 1998.

26. Sean R Eddy. Accelerated profile HMM searches. PLoS Comput Biol, 7(10):e1002195, 2011.

27. Jack W Roddy, David H Rich, and Travis J Wheeler. nail: software for high-speed, high-sensitivity protein sequence annotation. bioRxiv, 2024.

28. Kevin Karplus, Christian Barrett, and Richard Hughey. Hidden Markov models for detecting remote protein homologies. Bioinformatics (Oxford, England), 14(10):846–856, 1998.

29. Christiam Camacho, George Coulouris, Vahram Avagyan, Ning Ma, Jason Papadopoulos, Kevin Bealer, and Thomas L Madden. BLAST+: architecture and applications. BMC bioinformatics, 10:1–9, 2009.

30. Szymon M Kielbasa, Raymond Wan, Kengo Sato, Paul Horton, and Martin C Frith. Adaptive seeds tame genomic sequence comparison. Genome research, 21(3):487–493, 2011.

31. Michael Gribskov, Andrew D McLachlan, and David Eisenberg. Profile analysis: Detection of distantly related proteins. Proceedings of the National Academy of Sciences, 84(13):4355–4358, 1987.

32. Lawrence R Rabiner. A tutorial on hidden markov models and selected applications in speech recognition. Proceedings of the IEEE, 77(2):257–286, 1989.

33. Tim Anderson and Travis Wheeler. An FPGA-based hardware accelerator supporting sensitive sequence homology filtering with profile hidden markov models. bioRxiv, 2023.

34. Qi Chen, Bing Zhao, Haidong Wang, Mingqin Li, Chuanjie Liu, Zengzhong Li, Mao Yang, and Jingdong Wang. SPANN: Highly-efficient billion-scale approximate nearest neighbor search. In 35th Conference on Neural Information Processing Systems (NeurIPS 2021), 2021.

35. Cong Fu, Chao Xiang, Changxu Wang, and Deng Cai. Fast approximate nearest neighbor search with the navigating spreading-out graph. arXiv preprint, 2017.

36. Ruiqi Guo, Philip Sun, Erik Lindgren, Quan Geng, David Simcha, Felix Chern, and Sanjiv Kumar. Accelerating large-scale inference with anisotropic vector quantization. In International Conference on Machine Learning, pages 3887–3896. PMLR, 2020.

37. Yu A Malkov and Dmitry A Yashunin. Efficient and robust approximate nearest neighbor search using hierarchical navigable small world graphs. IEEE transactions on pattern analysis and machine intelligence, 42(4):824–836, 2018.

38. Masajiro Iwasaki and Daisuke Miyazaki. Optimization of indexing based on k-nearest neighbor graph for proximity search in high-dimensional data. arXiv preprint, 2018.

39. Jeff Johnson, Matthijs Douze, and Hervé Jégou. Billion-scale similarity search with GPUs. IEEE Transactions on Big Data, 7(3):535–547, 2019.

40. Kaiming He, Xiangyu Zhang, Shaoqing Ren, and Jian Sun. Deep residual learning for image recognition. In Proceedings of the IEEE conference on computer vision and pattern recognition, pages 770–778, 2016.

41. Djork-Arné Clevert Thomas Unterthiner, and Sepp Hochreiter. Fast and accurate deep network learning by exponential linear units (elus). arxiv 2015. arXiv preprint arxiv:1511.07289, 2020.

42. Kihyuk Sohn. Improved deep metric learning with multi-class N-pair loss objective. Advances in neural information processing systems, 29, 2016.

43. Martin C Frith. A new repeat-masking method enables specific detection of homologous sequences. Nucleic acids research, 39(4):e23–e23, 2011.

44. Daniel R Olson and Travis J Wheeler. Ultra-effective labeling of tandem repeats in genomic sequence. Bioinformatics Advances, 4(1):vbae149, 2024.

45. Diederik P. Kingma and Jimmy Ba. Adam: A method for stochastic optimization, 2017.

46. Martin C Frith, Michiaki Hamada, and Paul Horton. Parameters for accurate genome alignment. BMC bioinformatics, 11:1–14, 2010.

47. Milot Mirdita, Lars Von Den Driesch, Clovis Galiez, Maria J Martin, Johannes Söding, and Martin Steinegger. Uniclust databases of clustered and deeply annotated protein sequences and alignments. Nucleic acids research, 45(D1):D170–D176, 2017.

48. Baris E Suzek, Yuqi Wang, Hongzhan Huang, Peter B McGarvey, Cathy H Wu, and UniProt Consortium. UniRef clusters: a comprehensive and scalable alternative for improving sequence similarity searches. Bioinformatics, 31(6):926–932, 2015.

49. Tymor Hamamsy, James T Morton, Robert Blackwell, Daniel Berenberg, Nicholas Carriero, Vladimir Gligorijevic, Charlie EM Strauss, Julia Koehler Leman, Kyunghyun Cho, and Richard Bonneau. Protein remote homology detection and structural alignment using deep learning. Nature biotechnology, 42(6):975–985, 2024.

50. Wei Liu, Ziye Wang, Ronghui You, Chenghan Xie, Hong Wei, Yi Xiong, Jianyi Yang, and Shanfeng Zhu. Plmsearch: Protein language model powers accurate and fast sequence search for remote homology. Nature communications, 15(1):2775, 2024.

51. Zeming Lin, Halil Akin, Roshan Rao, Brian Hie, Zhongkai Zhu, Wenting Lu, Nikita Smetanin, Robert Verkuil, Ori Kabeli, Yaniv Shmueli, et al. Evolutionary-scale prediction of atomic-level protein structure with a language model. Science, 379(6637):1123–1130, 2023.

52. Maxat Kulmanov, Francisco J Guzmán-Vega, Paula Duek Roggli, Lydie Lane, Stefan T Arold, and Robert Hoehndorf. Protein function prediction as approximate semantic entailment. Nature Machine Intelligence, 6(2):220–228, 2024.

53. Benjamin Giovanni Iovino, Haixu Tang, and Yuzhen Ye. Protein domain embeddings for fast and accurate similarity search. In International Conference on Research in Computational Molecular Biology, pages 421–424. Springer, 2024.

54. Konstantin Schütze, Michael Heinzinger, Martin Steinegger, and Burkhard Rost. Nearest neighbor search on embeddings rapidly identifies distant protein relations. Frontiers in Bioinformatics, 2:1033775, 2022.

55. Yang Zhang and Jeffrey Skolnick. Scoring function for automated assessment of protein structure template quality. Proteins: Structure, Function, and Bioinformatics, 57(4):702–710, 2004.

56. David Y. Hancock, Jeremy Fischer, John Michael Lowe, Winona Snapp-Childs, Marlon Pierce, Suresh Marru, J. Eric Coulter, Matthew Vaughn, Brian Beck, Nirav Merchant, Edwin Skidmore, and Gwen Jacobs. Jetstream2: Accelerating cloud computing via jetstream. In Practice and Experience in Advanced Research Computing 2021: Evolution Across All Dimensions, PEARC ‘21, New York, NY, USA, 2021. Association for Computing Machinery.

57. Timothy J. Boerner, Stephen Deems, Thomas R. Furlani, Shelley L. Knuth, and John Towns. Access: Advancing innovation: NSF’s advanced cyberinfrastructure coordination ecosystem: Services & support. In Practice and Experience in Advanced Research Computing 2023: Computing for the Common Good, PEARC ‘23, page 173–176, New York, NY, USA, 2023. Association for Computing Machinery.

58. Claire D McWhite, Isabel Armour-Garb, and Mona Singh. Leveraging protein language models for accurate multiple sequence alignments. Genome Research, 33(7):1145–1153, 2023.

